# Discovery of an autoimmunity-associated *IL2RA* enhancer by unbiased targeting of transcriptional activation

**DOI:** 10.1101/091843

**Authors:** Dimitre R. Simeonov, Benjamin G. Gowen, Mandy Boontanrart, Theodore Roth, Youjin Lee, Alice Chan, Michelle L. Nguyen, Rachel E. Gate, Meena Subramaniam, Jonathan M. Woo, Therese Mitros, Graham J. Ray, Nicolas L. Bray, Gemma L. Curie, Nicki Naddaf, Eric Boyer, Frederic Van Gool, Kathrin Schumann, Mark J. Daly, Kyle K. Fahr, Chun Ye, Jeffrey A. Bluestone, Mark S. Anderson, Jacob E. Corn, Alexander Marson

## Abstract

The majority of genetic variants associated with common human diseases map to enhancers, non-coding elements that shape cell type-specific transcriptional programs and responses to specific extracellular cues ^1-3^. In order to understand the mechanisms by which non-coding genetic variation contributes to disease, systematic mapping of functional enhancers and their biological contexts is required. Here, we develop an unbiased discovery platform that can identify enhancers for a target gene without prior knowledge of their native functional context. We used tiled CRISPR activation (CRISPRa) to synthetically recruit transcription factors to sites across large genomic regions (>100 kilobases) surrounding two key autoimmunity risk loci, *CD69* and *IL2RA* (interleukin-2 receptor alpha; *CD25*). We identified several CRISPRa responsive elements (CaREs) with stimulation-dependent enhancer activity, including an *IL2RA* enhancer that harbors an autoimmunity risk variant. Using engineered mouse models and genome editing of human primary T cells, we found that sequence perturbation of the disease-associated *IL2RA* enhancer does not block *IL2RA* expression, but rather delays the timing of gene activation in response to specific extracellular signals. This work develops an approach to rapidly identify functional enhancers within non-coding regions, decodes a key human autoimmunity association, and suggests a general mechanism by which genetic variation can cause immune dysfunction.

---

Systematic studies of enhancer function remain challenging because of our limited understanding of the cellular contexts where each enhancer contributes to gene regulation. Functional enhancers can be mapped using Cas9-directed mutagenesis to disrupt genomic sequences^4-6^, but this approach only identifies the subset of enhancers that are necessary in the particular cellular context being studied. We hypothesized that localization of a strong transcriptional activator to an enhancer would be sufficient to drive target gene expression via transcription factor recruitment. This approach should be independent of the physiological cellular context in which the enhancer functions, and could “fire” an enhancer that has been poised by existing chromatin state. There has been recent success in engineering CRISPR activation (CRISPRa) systems by fusing nuclease-dead Cas9 (dCas9) to transcriptional activation domains such as VP64 (dCas9-VP64)^7-11^. CRISPRa is typically used to activate genes by targeting their promoters, but recent work suggests that CRISPRa can also transactivate target genes from distal enhancers^10,12^. We adopted this approach for high-throughput functional enhancer discovery with large libraries of guide RNAs (gRNAs) that tile genomic loci of interest (Fig. 1a).

**Figure 1.**
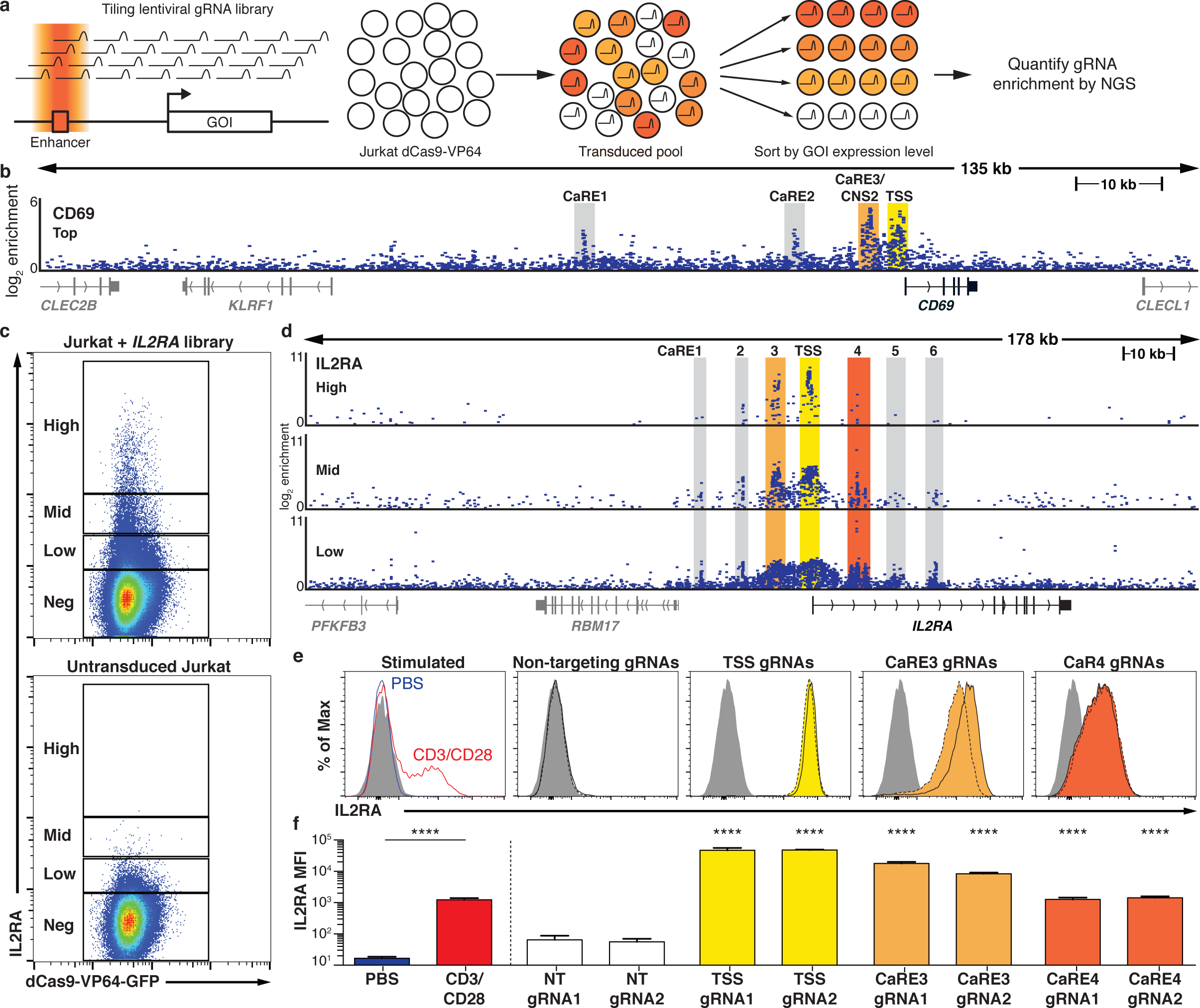
Discovery of putative enhancers with a tiling CRISPRa screen. (a) Schematic of the CRISPRa screen workflow. (b) Genomic coordinates of gRNAs plotted against enrichment into the "CD69 Top” sorted population. Each point represents an individual gRNA, plotted as fold-enrichment over unsorted cells. Peaks of guide activity are highlighted. (c) Distribution of IL2RA expression on Jurkat-dCas9VP64 cells transduced with the IL2RA tiling gRNA library. (d) Genomic coordinates of gRNAs plotted against enrichment into the IL2RA "High”, "Mid”, and "Low” sorted population, plotted as in (b). (e) Representative flow cytometry plots of IL2RA expression on Jurkat cells transduced with dCas9-VP64 and individual gRNAs. For each target region, solid black lines represent gRNA 1 and dashed black lines represent gRNA 2. Shaded gray histograms represent isotype control staining. Cells stimulated for 48 h with plate-bound CD3 and CD28 antibodies are shown for comparison. (f) Isotype-subtracted geometric mean fluorescence intensity (MFI) of data in (e). Data are presented as mean ± s.d., n=3 biological replicates. Statistical tests were performed on log-transformed MFI values. PBS control and anti-CD3/CD28 stimulated samples were compared using an unpaired two-tailed student’s t test. TSS and CaRE gRNA samples were compared to each non-targeting (NT) gRNA using one-way ANOVA followed by Sidak’s multiple comparisons test. Data in (e) and (f) are representative of at least 2 independent experiments. **** p<0.0001.

We tested the ability of tiled CRISPRa to reveal functional enhancers at the *CD69* locus, which contains multiple autoimmunity risk associations and a previously characterized stimulation-responsive enhancer^13^. CD69 is a cell surface receptor that is rapidly induced on T cells in response to T cell receptor (TCR) stimulation^14^. We sought to determine if CRISPRa could identify *CD69* cis-regulatory elements in resting cells, even in the absence of TCR stimulation. We transduced resting Jurkat cells stably expressing dCas9-VP64 with a pooled lentiviral guide RNA library that targeted all sites adjacent to an *S. pyogenes* Cas9 PAM 100 kb upstream of the *CD69* transcription start site (TSS), through the gene body, and 25 kb downstream of *CD69.* In total, the *CD69* library comprised 10,780 gRNAs and covered a 135 kb window. Transduction was performed at a low multiplicity of infection to ensure that only one gRNA was present per cell. We sorted transduced cells into four bins of CD69 expression and measured the distribution of gRNAs in the sorted populations (Fig. 1b, Supplementary Fig. 1). As expected, the top CD69 expressing cells were enriched for gRNAs targeting the *CD69* transcriptional start site (TSS) (Fig. 1b)^8^. We also observed enrichment for gRNAs well outside the canonical CRISPRa window. One of these CRISPRa Responsive Elements (CaREs) represents the previously characterized stimulation-responsive enhancer referred to as conserved non-coding sequence 2 (CNS2)^13^. Hence, tiling a transcriptional activator (dCas9-VP64) to non-coding sequences can identify enhancers independent of their functional context.

We next applied our approach to the *IL2RA* locus. *IL2RA*, also known as CD25, encodes a subunit of the high affinity interleukin-2 (IL2) receptor. Genome-wide association studies (GWAS) have implicated non-coding variants in the *IL2RA* locus as risk factors for at least eight autoimmune disorders, underscoring the critical role of *IL2RA* regulation in human immune homeostasis^1^. However, the functional impact of disease variants remains unclear because of the complex regulatory landscape at the *IL2RA* locus that is responsive to multiple signals. In resting conventional T cells, *IL2RA* is not only induced by antigen stimulation via the TCR, but is also potently regulated by a number of other signals. Regulators of *IL2RA* expression include cytokines like IL2, which upregulates the receptor as part of a positive feedback loop^15,16^. Furthermore, *IL2RA* regulation is also dependent on cellular programming. FOXP3+ regulatory T cells (Tregs), which are required to suppress auto-reactive T cells and prevent the development of multi-organ autoimmunity, constitutively express high levels of IL2RA and depend on it for their survival^17^. We hypothesized that multiple extracellular and cell-type specific signals are integrated to regulate gene expression through effects on distinct enhancer elements within the T cell super-enhancer at the *IL2RA* locus^18,19^. Whereas coding mutations in the gene affect all cell types that express IL2RA^20^, disease-associated non-coding variants could selectively affect *IL2RA* induction in conventional T cells in response to a specific signal or impair constitutive expression in Tregs. We sought to map functional *IL2RA* enhancer elements and determine how known disease risk variants affect enhancer function.

To discover *IL2RA* CaREs, we transduced Jurkat-dCas9-VP64 cells with a library of 20,412 gRNAs tiling 178 kb around the *IL2RA* locus. This caused a range of increased IL2RA expression on the transduced cells(Fig. 1c). Transduced cells were sorted into four bins of increasing expression (“Negative”, “Low”, “Mid”, and “High”). Analysis of gRNAs enriched in each expression bin revealed at least six CaREs leading to different levels of IL2RA expression: three in the first intron and three upstream of the promoter (Fig. 1d). We confirmed that individual gRNAs targeting *IL2RA* CaRE3 and CaRE4 transactivate IL2RA in both unstimulated Jurkat and HuT78 human T cell lines to levels comparable to those resulting from activation (Fig. 1e-f, Supplementary Fig. 2). The expression level of IL2RA induced by individual gRNAs was correlated with enrichment of those gRNAs during library sorting (Supplementary Fig. 3). In sum, our unbiased transcriptional activation approach identified novel elements within the *IL2RA* super-enhancer where transcription factor recruitment is sufficient to induce IL2RA expression on resting cells.

**Figure 2.**
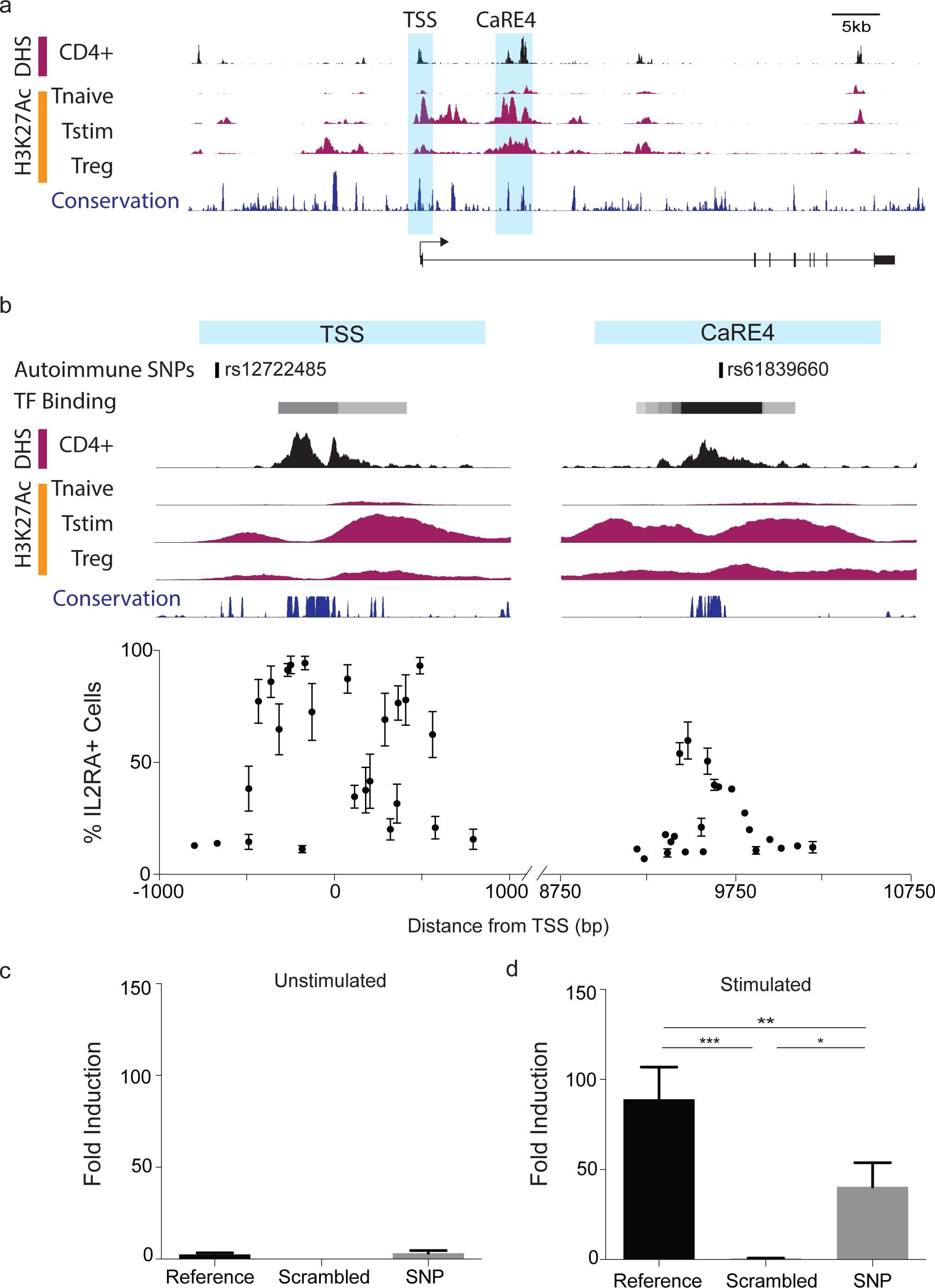
Identification of a stimulation-dependent disease-associated IL2RA enhancer. (a) The IL2RA locus overlapped with DNAse hypersensitive sites (DHS), H3K27Ac from primary human T cells (Epigenome Roadmap) and vertebrate conservation (PhastCons 46-way) highlighting the genomic features at the IL2RA transcriptional start site (TSS) and CaRE4. (b) Zoomed in view of the IL2RA TSS and a region of CaRE4 marked by an autoimmunity SNP showing TF binding (ENCODE), H3K27Ac and conservation as in (a). Jurkat-dCas9-VP64 cells were nucleofected with gRNA expression plasmids. IL2RA expression on nucleofected (BFP+) cells was analyzed by flow cytometry 48 hours post-nucleofection. Data in (b) are presented as mean +/- s.d. (n=2) and are pooled from 2 independent experiments. (c) Jurkat cells were nucleofected with luciferase reporter constructs containing a minimal promoter downstream of the CaRE4 reference sequence, a scrambled sequence, or CaRE4 with rs61839660 (SNP). Cells were lysed and luciferase activity was measured 1 day post-nucleofection. (d) Jurkat cells nucleofected as in (c). Cells were stimulated with plate bound anti-CD3 and anti-CD28 antibodies 1 day post-nucleofection. Cells were lysed and luciferase activity was measured 20 hours post-stimulation. Data in (c) and (d) are presented as mean +/- s.d. and are representative of at least two independent experiments. A one-way ANOVA (significance level = 0.05) followed by Holm Sidak’s multiple comparisons test was used statistical analysis comparing scrambled and SNP enhancer sequences to reference sequence in the luciferase assays. *p<0.05, **p<0.01, and ***p<0.001.

Our results at the *CD69* locus demonstrated that CaREs can represent enhancers^13^ and we next tested whether the *IL2RA* CaREs we identified are *bona fide* T cell enhancers. Active enhancers often have stereotyped genomic characteristics, including evolutionary sequence conservation, chromatin accessibility, and signature histone modifications, such as H3K27 acetylation (H3K27Ac). CaRE4 and several other *IL2RA* CaREs lack H3K27Ac in IL2RA-negative naïve T cells but exhibit increased H3K27Ac in IL2RA-positive stimulated T cells and Tregs^1^, consistent with their possible function as context-dependent enhancers (Fig. 2a). We initially defined CaREs using broad windows, but we next tested gRNAs in an arrayed fashion to determine if the ones that caused the strongest induction of IL2RA overlap with canonical enhancer characteristics(Fig. 2b). We expressed individual gRNAs surrounding the TSS and within CaRE4 to more finely map the genomic features of sites where CRISPRa can transactivate *IL2RA.* The strongest transactivation in CaRE4 was observed when dCas9-VP64 was recruited to a highly conserved nucleosome-depleted region bound by multiple TFs between two peaks of H3K27ac in stimulated T cells. These sites of peak CRISPRa activity also overlapped with conserved sequences marked by chromatin accessibility in primary CD4+ T cells in DNase hypersensitivity assays^2,21^. Finally, to experimentally test the enhancer activity of CaRE4 we cloned ~500 bases of the element (centered on the conserved sequence) into a luciferase reporter vector upstream of a minimal promoter. Notably, this region of CaRE4 drove strong luciferase expression in Jurkat T cells, but only in response to stimulation (Fig. 2c-d), validating *IL2RA* CaRE4 as a stimulation-dependent enhancer.

Sequence variation in CaRE4 has been implicated in risk of human autoimmunity (Fig. 2b). The single nucleotide polymorphism (SNP) rs61839660, which resides in this element, is one of only a few disease-associated variants for any common human disease that has been statistically resolved convincingly to a single non-coding variant. This individual SNP accounts for the risk of inflammatory bowel disease (IBD) at the *IL2RA* locus^22^. Consistent with a critical and complex function in immune regulation, this same SNP also paradoxically causes protection from type 1 diabetes (T1D) in the locus^23^. Having confirmed that CaRE4 is a stimulation dependent enhancer, we tested if rs61839660 affects enhancer function. Indeed, mutation of CaRE4 to introduce the autoimmune SNP was sufficient to blunt its stimulation-induced activity in the luciferase assay (Fig. 2c-d). These findings link rs61839660’s role in disease to disruption of a stimulation-dependent *IL2RA* enhancer.

The *IL2RA* CaRE4 enhancer is highly conserved between human and mouse, which allowed us to test the *in vivo* effects of sequence variation. We used Cas9 genome editing to generate knock-in mice with the human autoimmune-associated SNP or a 12 bp deletion (12DEL) (Fig. 3a). Founders were backcrossed and bred to homozygosity for immunophenotyping at 2-4 months of age. *In vivo* phenotyping of the enhancer-edited mice and littermate controls revealed no evidence of overt immune dysregulation (Supplementary Fig. 4). T cell development in these mice was normal with no differences in total thymic cell number or percentages of T cell developmental stages (Supplementary Fig. 5). Genetic alterations in the *IL2RA* enhancer had no consistent effect on the abundance of regulatory T cells, which constitutively express *IL2RA* (Supplementary Fig. 6). SNP and 12DEL Tregs also expressed normal levels of IL2RA, suggesting that the enhancer is not required for overall IL2RA expression on CD4+ T cells at steady state (Fig. 3b and Supplementary Fig. 6).

**Figure 3.**
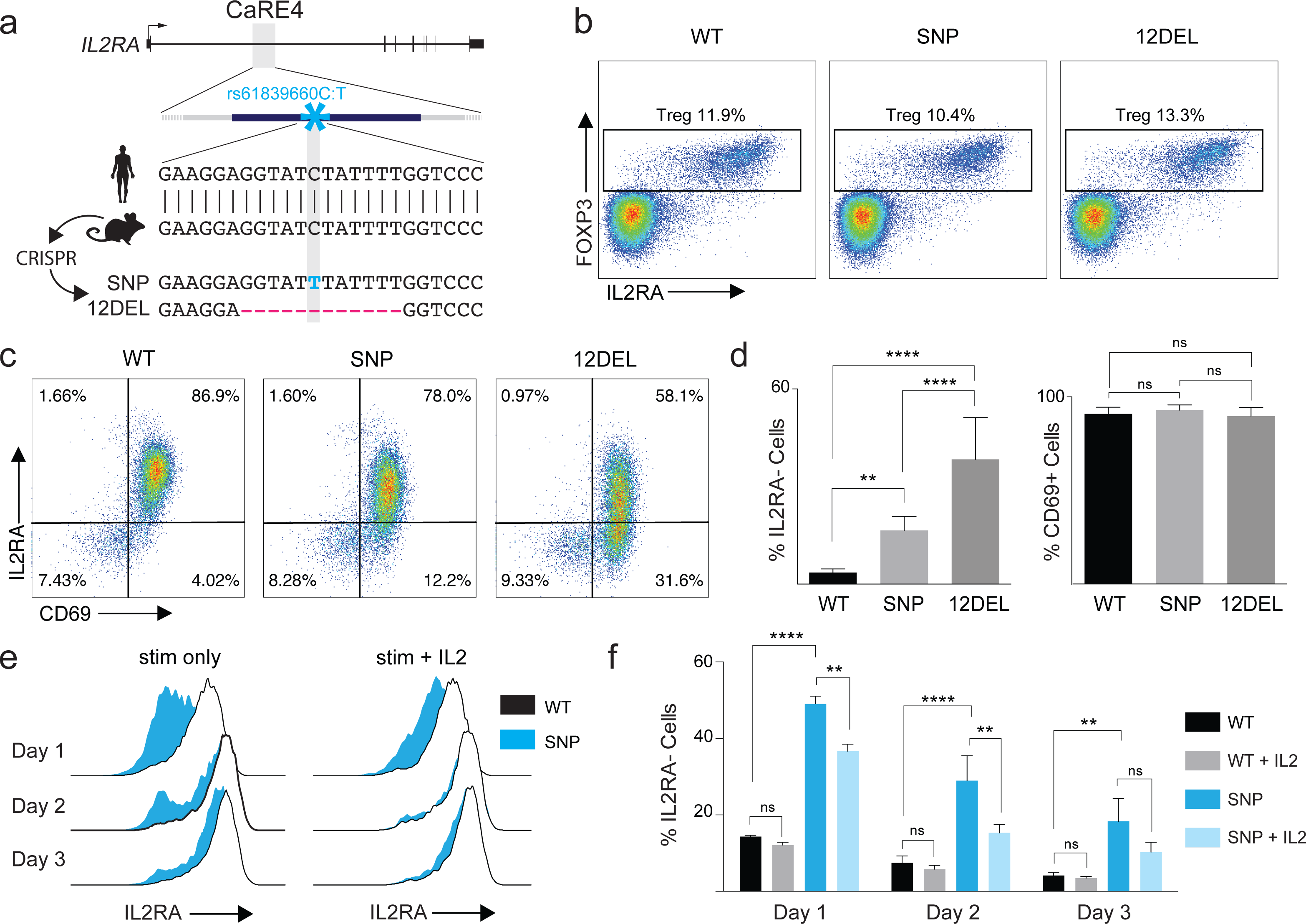
In vivo modeling of sequence variation in the IL2RA enhancer. (a) Generation of rs61839660 knock-in (SNP) and 12 bp deletion (12DEL) B6 mice using CRISPR to edit the conserved IL2RA enhancer in zygotes. (b) Peripheral lymph node staining of regulatory T cells (CD4+FOXP3+IL2RA+). (c) Representative flow cytometry plots for IL2RA and CD69 surface staining on naive T cells stimulated with plate-bound anti-CD3 and anti-CD28 antibodies for 1 day. (d) Quantification of the percent IL2RA- and CD69+ cells 1 day after stimulation. (e) 3-day time course of naive T cells stimulated with antibodies alone or in combination with 50U/ml IL2. (f) Quantification of flow cytometry data in panel (e). All data are presented as mean +/- s.d. and are representative of at least two independent experiments. Biological replicates of WT (n=7), SNP (n=4) and 12DEL (n=5) were used. A one-way ANOVA (statistical significance = 0.05) followed by Holm Sidak’s multiple comparisons test was used for stastical analysis comparing enhancer edited conditions to wildtype conditions.**p<0.01, ****p<0.001

Given the stimulation-dependent enhancer activity in human cells *in vitro*, we reasoned that the CaRE4 enhancer might regulate *IL2RA* induction on naïve CD4+ T cells following stimulation. We isolated naïve T cells (CD4+CD62L+CD44-) and activated them *in vitro* with anti-CD3 and anti-CD28 antibodies. Remarkably, naïve T cells from both SNP and 12DEL mice had significantly reduced IL2RA surface expression compared to wild type (WT) mice 24 hours post-activation (Fig. 3c and Fig. 3d). The deficit was more pronounced in 12DEL cells, but the SNP alone resulted in significant reduction of IL2RA levels (50% of WT IL2RA, p<0.0001 two-tailed t-test). Reduced IL2RA levels were not due to a general defect in response to stimulation, as CD69 expression was induced to levels comparable to WT cells after 24 hours of stimulation (Fig. 3d). Since disruption of CaRE4 did not ablate steady state IL2RA, we asked whether mutant T cells are able to recover IL2RA levels at longer time points after stimulation. Indeed, three days post-stimulation, IL2RA levels on naïve T cells from SNP mice recovered almost completely to wild type levels (Fig. 3e-f). We next asked if other extracellular signals could compensate for the disease variant’s effect on the timed response to T cell activation. We found that IL2 stimulation accelerated the rate at which T cells from SNP mice induced IL2RA on their surface. Our results suggest that the human rs61839660 disease-associated variant impairs the function of an intronic enhancer, and that this enhancer regulates IL2RA induction in response to activation of naïve T cells. Furthermore, this impairment can be alleviated by IL2 signaling. The disease-associated single nucleotide change has subtle effects on final levels of IL2RA, but exerts a pronounced effect on the timing of induction.

Having established a role for the CaRE4 enhancer and associated variants in mouse models, we asked whether the timing of IL2RA expression is similarly regulated in primary human T cells. We previously showed that Cas9 ribonucleoproteins (RNP) can be used to efficiently edit primary human T cell genomes and generate both knock-in and knock-out cells^24^. We used a cocktail of RNPs to target the enhancer, combining three gRNAs surrounding the autoimmunity SNP within CaRE4. We sorted primary naïve T cells (CD4+IL2RA-CD127+CD45RA+CD45RO-) from human blood and electroporated them with RNPs to introduce edits in the *IL2RA* locus (Fig. 4a). Electroporated cells were activated with anti-CD3/anti-CD28 antibodies and assessed for IL2RA cell surface expression over the course of three days. The targeted cells exhibited strongly impaired IL2RA upregulation at 24 hours (Fig. 4b), but by 72 hours expression levels almost matched those of cells treated with a non-targeting RNP. These results are consistent with our findings in edited murine cells. The IL2RA phenotype was linked to specific genotypes created at the CaRE4 enhancer by Cas9 mutagenesis. At 24 hours, IL2RA-cells showed an enrichment of large enhancer deletions, as detected by PCR product size (Fig. 4c). Furthermore, IL2RA-cells were enriched for insertion/deletion mutations around the rs61839660 SNP, as measured by deep amplicon sequencing (Supplementary Fig. 7).

**Figure 4.**
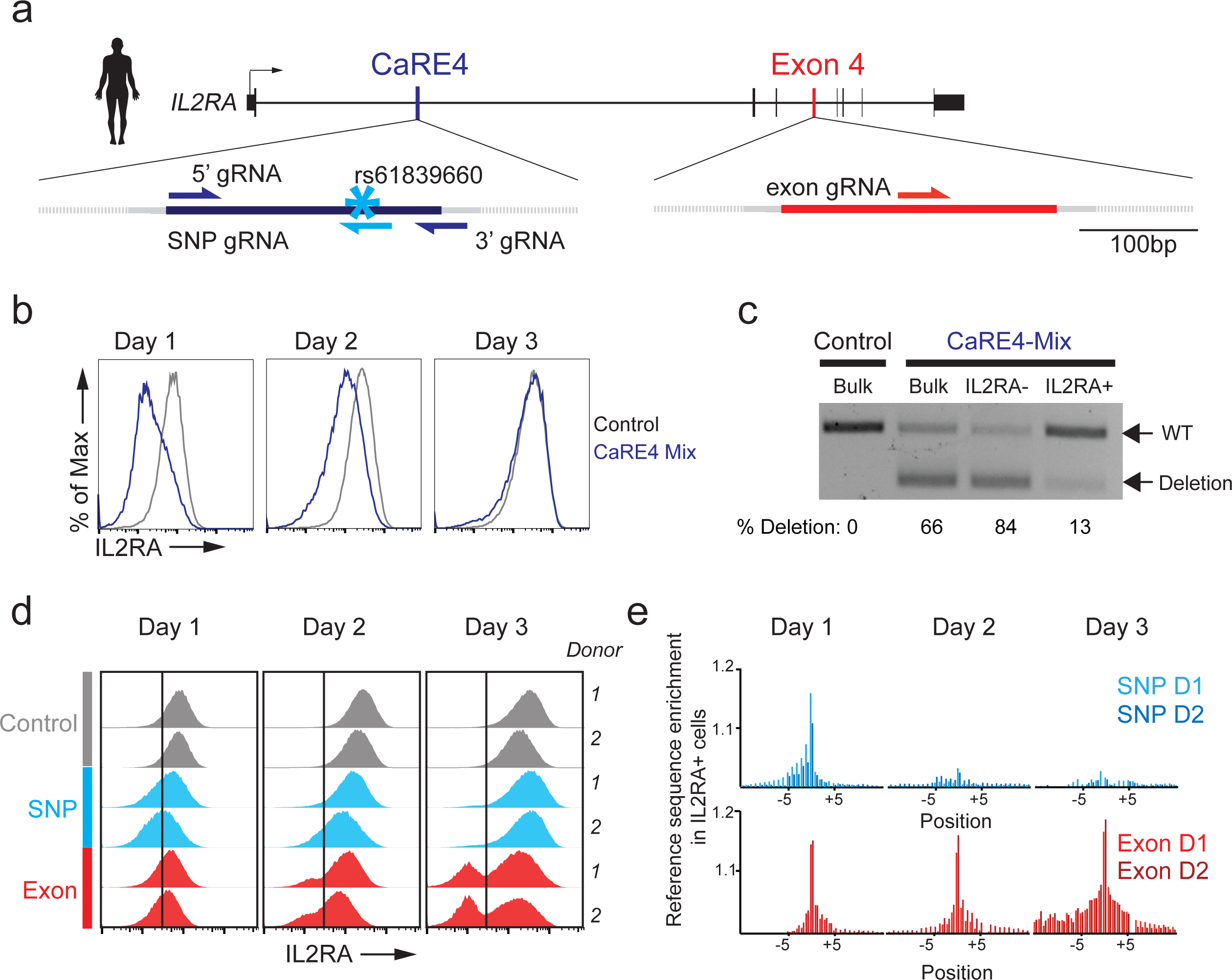
Enhancer-editing in primary human T cells impairs IL2RA kinetics. (a) Targeting strategy to edit the IL2RA enhancer and IL2RA coding sequences with Cas9 ribonucleoproteins (RNPs) in primary human T cells. (b) Naive T cells were electroporated with three RNPs (CaRE4 Mix) directed against the IL2RA enhancer or a non-targeting control gRNA (Control), and assessed for IL2RA expression over three days after stimulation. (c) PCR amplification of unsorted (bulk) or sorted IL2RA- and IL2RA+ enhancer targeted cells after 1 day of stimulation from (b) shows enrichment for a large enhancer deletion in the IL2RA-population. (d) To assess how sequence variation at the site of the SNP affects IL2RA induction, primary human naive T cells were electroporated with single RNP complexes against the SNP site, exon 4 of IL2RA or a non-targeting RNP. IL2RA expression was assessed in these cells by flow cytometry over three days of stimulation. (e) IL2RA+ cells from (d) were sorted every day and deep sequenced at the enhancer and exon 4 cut sites to analyze sequence edits. Enrichment of reference sequence in the IL2RA+ cells compared to unsorted cells is shown for both donors. Whereas enrichment of reference sequence is preserved in the population of IL2RA+ cells with coding sequence disruptions, IL2RA+ cells from non-coding targeting lose this enrichment with time.

We next sought to map functional nucleotides in CaRE4 at the site of the SNP. We electroporated cells with a single Cas9 RNP targeting a cut-site adjacent to the SNP and then FACS sorted IL2RA- and IL2RA+ enhancer-edited cells at 24, 48, and 72 hours post-stimulation (Fig. 4d). We used amplicon sequencing in each population to capture the insertion and deletion edits generated by Cas9 mutagenesis. By comparing the edits in the sorted populations to the background distribution of edits from unsorted edited cells, we were able to quantify the effects of specific sequences on enhancer function at each time point. On Day 1 post-stimulation, we observed that there was an enrichment of the unedited reference sequence around the cut site in the IL2RA+ population relative to the bulk unsorted cells, indicating that these sequences were critical for proper IL2RA expression at that time point (Fig. 4e). However, this preservation of the reference sequence was no longer evident by Day 3. In contrast, edits at a coding site in Exon 4 of *IL2RA* caused marked accumulation of IL2RA-cells, and the preservation of the reference sequence in the IL2RA+ population persisted through Day 3 (Fig. 4e). In sum, Cas9 RNP editing of primary human cells revealed the temporal effects of non-coding mutations and constitutive effects of coding mutations. An intact CaRE4 enhancer is essential to maintain normal timing of the human naïve T cell response to activation.

Here we show that CRISPRa is a powerful approach for high-throughput enhancer discovery that can be used to rapidly map functional enhancers without prior knowledge of their specific biological contexts. While we focused on immune-related genes, we anticipate this approach will have general utility as an enhancer discovery platform and can be used for annotation of the vast non-coding genomic space. This technology enabled us to discover a disease-associated enhancer that controls the timing of gene expression. rs61839660 is highly conserved across species, which allowed us to determine its function in both human and mouse primary T cells. However, the majority of non-coding disease variants diverge between mouse and human. Genome editing in human primary cells will be required to understand the function of non-coding sequence variation.

Determining the cellular contexts in which an enhancer regulates gene expression is critical to understanding how non-coding variation in the locus can contribute to pathology. We have identified an enhancer that tunes the kinetics of *IL2RA* gene activation in response to stimulation. Experimental ablation of non-coding elements recently suggested that a subset of enhancers have only temporary phenotypes, with target gene expression recovering over time^25^. Our findings now reveal that human non-coding disease variants can also cause temporally defined cellular phenotypes. Not only can autoimmunity risk variants shape response-dependent expression changes^34^, but the magnitude of genetically encoded responses can be temporally defined. Genetic variants such as rs61839660 may persist at high frequencies in the population because of tightly constrained temporal effects and only alter gene expression under specific conditions.

Our findings suggest that human immune homeostasis depends not only on normal IL2RA expression in Tregs, but also on proper kinetics of IL2RA induction in conventional T cells. Further study is needed to reveal the mechanistic link between altered IL2RA kinetics and autoimmunity, but IL2/IL2RA signaling is central to T cell proliferation, differentiation and survival. Ongoing clinical trials are testing the use of IL2 to treat various autoimmune and inflammatory conditions^26^. Understanding how genetics interact with IL2 signals to regulate IL2RA induction in may provide mechanistic insights relevant to such therapies and inform patient stratification decisions. The data presented here critically identifies a specific window where a genetic risk factor for autoimmunity acts, and our work defines a new model to understand how common genetic variants control cell-type specific temporal gene regulation in health and disease.

## Acknowledgements

We thank all members of Marson and Corn labs and Abul Abbas, Stanley Qi, Luke Gilbert, Jonathan Weissman, Marc Gavin and Warren Leonard for suggestions and technical assistance. This research was supported by the UCSF Sandler Fellowship (A.M.), a gift from Jake Aronov (A.M.) and National Multiple Sclerosis Society (CA 1074-A-21). Alexander Marson, M.D., Ph.D. holds a Career Award for Medical Scientists from the Burroughs Wellcome Fund. Benjamin Gowen is supported by the IGI-AstraZeneca Postdoctoral Fellowship. We thank Jackson Laboratories for generating the SNP mice and Agilent for generating oligos for cloning of the CRISPRa gRNA library. We thank the UC Berkeley High Throughput Screening Facility for preparation of gRNA lentivirus and the UC Berkeley Flow Cytometry Facility for cell sorting. This work used the Vincent J. Coates Genomics Sequencing Laboratory at UC Berkeley, supported by NIH S10 Instrumentation Grants S10RR029668 and S10RR027303. A patent has been filed on the use of Cas9 RNPs to edit the genome of human primary T cells (A.M.). A.M. serves as an advisor to Juno Therapeutics and the Marson lab has had sponsored research agreements with Juno Therapeutics and Epinomics.

## Figure Legends

**Supplementary Figure 1. Expression of CD69 on gRNA-expressing cells.** (a) Distribution of CD69 expression on Jurkat-dCas9-VP64 cells transduced with the CD69 tiling gRNA library. (b),(c) Representative flow cytometry plots of CD69 expression on Jurkat (b) or HuT78 cells (c) transduced with dCas9-VP64 and individual gRNAs. For each target region, solid black lines represent gRNA 1 and dashed black lines represent gRNA 2. Shaded gray histograms represent isotype control staining. Cells stimulated for 48 h with plate-bound CD3 and CD28 antibodies are shown for comparison. (d),(e) Isotype-subtracted geometric mean fluorescence intensity (MFI) of data in (b) and (c). Statistical tests were performed on log-transformed MFI values. PBS control and anti-CD3/CD28 stimulated samples (CD3/CD28) were compared using an unpaired two-tailed student’s t test. TSS and CaRE gRNA samples were compared to each nontargeting (NT) gRNA sample using one-way ANOVA followed by Sidak’s multiple comparisons test. Data are presented as mean ± s.d., n=3 biological replicates. Data are representative of at least 2 independent experiments. ** p<0.01, *** p<0.001, **** p<0.0001.

**Supplementary Figure 2. Upregulation of IL2RA expression by CaRE gRNAs in HuT78 cells.** (a) Representative flow cytometry plots of CD69 expression on HuT78 cells transduced with dCas9-VP64 and individual gRNAs. For each target region, solid black lines represent gRNA 1 and dashed black lines represent gRNA 2. Shaded gray histograms represent isotype control staining. Cells stimulated for 48 h with plate-bound CD3 and CD28 antibodies are shown for comparison. (b) Isotype-subtracted geometric mean fluorescence intensity (MFI) of data in (b). Statistical tests were performed on log-transformed MFI values. PBS control and antiCD3/CD28 stimulated samples (CD3/CD28) were compared using an unpaired two-tailed student’s t test. TSS and CaRE gRNA samples were compared to each non-targeting (NT) gRNA sample using one-way ANOVA followed by Sidak’s multiple comparisons test. Data are presented as mean ± s.d., n=3 biological replicates. Data are representative of at least 2 independent experiments. **** p<0.0001.

**Supplementary Figure 3. Effects of individual gRNAs are correlated with enrichment in the IL2RA screen.** Jurkat dCas9-VP64 cells were transduced with individual gRNAs from the IL2RA library, and surface IL2RA expression was measured by flow cytometry. The isotype-subtracted geometric mean fluorescence intensity (MFI) of the transduced cells is plotted against gRNA enrichment in the indicated IL2RA bin in the IL2RA screen.

**Supplementary Figure 4. IL2RA enhancer-edited mice show no evidence of immune dysfunction.** Enhancer-edited mice and littermate controls were immunophenotyped at 2–4 months of age. (a) Spleens from WT, SNP and 12DEL mice. (b) Total number of cells in spleen, peripheral lymph nodes (peri-LNs) and thymus. (c) Percentage of naive (CD4+CD62L+CD44-) and memory (CD4+CD62L-CD44+) cells in CD4 enriched T cells isolated from spleen and peri-LNs. All data are presented as mean +/- s.d. and are representative of at least two independent experiments. Data are biological replicates of WT (n=7), SNP (n=4), and 12DEL (n=5) mice. A non-parametric one-way ANOVA (significance level = 0.05) followed by Dunn’s multiple comparison test was used for statistical analysis comparing enhancer-edited mice to WT controls.

**Supplementary Figure 5. Normal T cell development in CaRE4 edited mice.** (a) Percentage of thymocytes in T cell developmental stages from the thymus of enhancer-edited mice and littermate controls at 2–4 months of age. Data are shown for CD4/CD8 single-positive (SP), double-positive (DP), and double-negative (DN) populations. (b) Quantification of IL2RA surface staining (geometric mean fluorescence intensity, MFI) on DN thymocytes. All data are presented as mean +/- s.d. and are representative of at least two independent experiments. Data are biological replicates of WT (n=7), SNP (n=4), and 12DEL (n=5) mice. A non-parametric one-way ANOVA (significance level = 0.05) followed by Dunn’s multiple comparison test was used for statistical analysis comparing enhancer edited mice to WT controls.

**Supplementary Figure 6. IL2RA enhancer-edited mice show no consistent regulatory T cell defects.** (a) Percent Tregs of CD4+ T cells in tissues of enhancer edited mice and littermate controls at 2–4 months of age. (b) Quantification of IL2RA surface staining (geometric mean fluorescence intensity, MFI) on FOXP3+ cells. All data are presented as mean +/- s.d. and are representative of at least two independent experiments. Data are biological replicates of WT (n=6), SNP (n=4), and 12DEL (n=5) mice. A non-parametric one-way ANOVA (significance level 0.05) followed by Dunn’s multiple comparison test was used to compare enhancer-edited mice to WT controls.*p<.05, **p<.01

**Supplementary Figure 7. IL2RA enhancer edits are linked to impaired IL2RA induction.** (a) Read pileups from deep sequencing at the IL2RA enhancer in CaRE4-mix-edited cells at 1 day after activation. The cut sites for the SNP gRNA and the 3’ gRNA are indicated with black arrows. The cut site for the 5’ gRNA was not captured here. Reference bases in the reads are shown in white and deviation from the reference is shown in color; deletions are colored red and insertions are colored blue. Editing enrichment is observed in the sorted IL2RA-cells as compared to the IL2RA+ cells from both donors.

**Supplemental Figure 8. Correlation of results across CRISPRa screen replicates.** Normalized read counts for gRNAs in the indicated cell populations are compared between biological replicates of the CD69 screen (a) and IL2RA screen (b).

**Supplementary Figure 9. Tracking rs61839660 knock-in alleles in isogenic primary human T cells links variant to impaired IL2RA induction.** (a) rs61839660 knock-in alleles were generated with a HDR template that was electroporated into the human primary T cells along with the RNPs. Deep sequencing was then carried out on unsorted cells as well as sorted IL2RA- and IL2RA+ cells over the course of the three days. Quantification of SNP knock-in reads as a percent of total reads in bulk and sorted populations is shown from two different donors.

## Materials and Methods

### Cell culture

Cell culture was performed at 37 C in a humidified atmosphere containing 5% CO2. Jurkat cells (Clone E6–1) were obtained from the Berkeley Cell Culture Facility. HuT78 cells were a gift from Art Weiss (UCSF, San Francisco, CA). Jurkat and HuT78 cells were cultured is RPMI1640 medium (Gibco) supplemented with 10% fetal bovine serum (FBS), 100 U/ml penicillin (Gibco), 100 μg/ml streptomycin (Gibco), and 1 mM sodium pyruvate (Gibco). Cell line identity was authenticated by STR analysis and verified mycoplasma free using the MycoAlert Mycoplasma Detection Kit (Lonza).

### Generation of dCas9-VP64 cells

Jurkat and HuT78 cells were transduced with a lentiviral dCas9-VP64–2A-GFP expression vector (Addgene 61422). Single GFP+ cells were sorted by FACS into the wells of a 96-well plate, and clones with bright uniform GFP expression were selected for use in future experiments.

### Antibodies

All antibodies used for staining or cell stimulation are listed in Supplemental Table 1.

### Tiling gRNA library generation

For each gene of interest, the window of tiling gRNA libraries extended from 100 kb upstream of the transcription start site through 25 kb downstream of the end of the gene. The hg19 coordinates of the *CD69* library window were chr12:9,880,082–10,013,497. The hg19 coordinates of the *IL2RA* library window were chr10:6,027,657-6,204,333.

gRNAs were designed against all NGG PAMs in the window, excluding sequences containing BstXI or BlpI/Bpu1102l cut sites. Each library contained 2,244 negative control gRNAs taken from the genome-scale CRISPRi/a libraries described in Gilbert et al^1^. Protospacer sequences flanked by restriction enzyme sites and PCR adaptors were synthesized by as pooled oligonucleotides by Agilent Technologies (Santa Clara, CA). Pooled gRNA libraries were then cloned into the lentiviral expression vector “pCRISPRia-v2” as described in Horlbeck et al^2^.

### Tiling transcriptional activation screen

Protocols for the pooled lentiviral CRISPRa screens were adapted from Gilbert et al^1^. Lentivirus was produced by transfecting HEK293T with standard packaging vectors using TransIT-LTI Transfection Reagent (Mirus, MIR 2306). Viral supernatant was harvested 48–72 hr following transfection, filtered through a 0.45 μm PES syringe filter, snap-frozen, and stored at -80 °C for future use.

Jurkat-dCas9-VP64 cells were infected with lentiviral gRNA libraries by resuspending cells at 2 × 10^6^ cells/mL in fresh media containing titered lentivirus and 4 μg/mL polybrene. Cells were spin-infected for 2 hours at 1000 xg, 33 °C, followed by resuspension in fresh media at 0.250.5 × 10^6^ cells/mL. To limit the number of cells expressing multiple gRNAs, lentivirus was titered to infect only 10–20% of cells. Cells were cultured in media containing 0.75 or 1.5 μg/mL puromycin for days 2–5 post-infection to remove uninfected cells. The number of initially infected cells was at least 500x the number of gRNAs in the library, and at least this many cells were maintained throughout the course of the experiment.

7–10 days post-infection cells were sorted based on IL2RA/CD25 or CD69 expression. Briefly, cells were resuspended in sterile sort buffer (PBS + 2% FBS) containing either CD25-PE or CD69-PE antibody at a 1:25 dilution. Cells were stained for 30 minutes on ice, washed twice with sort buffer, and passed through a 70 μm mesh. Cells were sorted into 4 bins based on IL2RA/CD25 or CD69 expression using a BD Influx cell sorter. The total number of cells collected was at least 500x the number of gRNAs in the library. Additional unsorted cells totaling 500x the number of gRNAs in the library were collected at this time. Duplicate infections and sorts were performed for each library. Collected cells were centrifuged at 500 xg for 5 minutes, and cell pellets were stored at -80 °C until genomic DNA was isolated.

Genomic DNA was isolated from sorted cells using NucleoSpin Blood kits (Macherey-Nagel), or by Proteinase K digestion and isopropanol precipitation for samples with fewer than 10^6^ cells. PCR was used to amplify gRNA cassettes with Illumina sequencing adapters and indexes as described in Kampmann, et al.^3^. Genomic DNA samples containing less than 10 µg of gDNA were loaded directly into PCR. For genomic DNA samples containing more than 10 µg of DNA, samples were first digested for 18 hours with SbfI-HF (NEB) to liberate a ~500 bp fragment containing the gRNA cassette. The gRNA cassette was isolated by gel electrophoresis as described in Kampmann, et al.^3^, and the isolated DNA was then used for PCR. Custom PCR primers are listed in Supplementary Table 2. Indexed samples were pooled and sequenced on an Illumina HiSeq-2500 with the custom sequencing primer 5’gtgtgttttgagactataagtatcccttggagaaccaccttgttg-3 ’. Sequencing libraries were pooled proportional to the number of sorted cells in each sample. The target sequencing depth was 2,000 reads/gRNA in the library for unsorted “background” samples, and 10 reads/cell in sorted samples.

### Screen data analysis

Sequence files were processed to remove low quality reads and reads lacking the gRNA constant region. Reads were then trimmed for the common sequence using the cutadapt script and the command “cutadapt -a GTTTAAGAGCTAAGCTG”^4^. Trimmed reads were then aligned against a database of the guide sequences using bowtie2 with option -norc^5^. Read counts for each sample were normalized to the total number of gRNA read counts in that sample. A pseudo-count of 1 was added to all normalized guide counts. gRNA enrichment was calculated as follows:

mean(log2(IL2RA_gate/IL2RA_background))

gRNA enrichment in sorted cell populations was visualized with the Integrative Genomics Viewer (Broad Institute). Non-targeting control gRNAs and gRNAs that map perfectly to multiple sequences within the gRNA library window were excluded from visualization. For each sorted cell population, normalized read counts for a given gRNA were well correlated between the two replicates of the screen (Supplementary Fig. 8).

### Screen validation

For screen validation using individual gRNAs, gRNAs were cloned into the same expression plasmid used for the gRNA library. Lentivirus was produced as described above and used to infect Jurkat and HuT78 cells expressing dCas9-VP64. Expression of IL2RA and CD69 on infected cells was analyzed by flow cytometry. A complete list of gRNAs used in CRISPRa follow-up experiments is provided in Supplementary Table 3.

### Transient gRNA expression

For the transient gRNA expression experiment shown in Fig. 2a, gRNAs were cloned into the same expression plasmid used for the gRNA library. 2 × 10^5^ Jurkat-dCas9-VP64 cells were nucleofected with 1 μg of sgRNA plasmid with a 4-D Nucleofector (Lonza) using 20 μL of Nucleofector Buffer SE and nucleofection program CL-120. CD25/IL2RA expression on nucleofected cells was analyzed by flow cytometry 48 hours post-nucleofection.

### Luciferase Assays

A 500 bp fragment centered on the *IL2RA* CaRE4 reference sequence was synthesized as a gBlock Gene Fragment (IDT, Coralville, IA, USA) and cloned into the Firefly Luciferase (Fluc) reporter vector pGL4.23 (Promega, Madison, WI, USA), upstream of a minimal promoter. Similarly we generated *IL2RA* CaRE4 constructs with the rs61839660C>T variant or with the conserved core of 350 bp scrambled. Each FLuc construct (700 ng) was electroporated with a Renilla luciferase plasmid (pGL4.74, 70 ng) into 5 × 10^5^ Jurkat cells using the 4-D Nucleofector, 20 μL Nucleofection buffer SE and nucleofection program CL-120. Cells were rested overnight and then activated using plate bound anti-CD3 (clone UCHT1, 10 μg/ml, TONBO Biosciences) and anti-CD28 (clone CD28.2, 10 μg/ml, TONBO Biosciences) antibodies for 22 hours. Luciferase expression was assessed using the Dual-Glo Luciferase Assay (Promega, Madison, WI, USA) on a 96 well plate luminometer. FLuc activity was normalized to Renilla activity and is reported as fold induction over empty pGL4.23 vector. The CaRE4 sequence fragments are listed below.

## >CaRE4-Reference

CCTACTGGCCGGTACCTGAGCTCGCTAGCCTCCCTCTTCCTCCTGTACCCTCAACCCATCAGCTGCCTGTACTGCTGTAGCAACAACTTCTCTCTGAAAACCTTCCCAGGGGAGAAAACGCGCCCCCCCTCACGCCTCAGAGGCACACACCTATCCTAGCCTTTGTTTTTTTATTGCTGAGAGTACAGAAAGCAGCGGCTTCTGAAGGAGGTATCTATTTTGGTCCCAAAC AGAAAAGAGT GGGT GAT TCTGT GGGC AGAGGT GGC AATT CCT GAAT GT GGT AT GGTGACTCACGCCCGGAGGTGCTCAGAATTTCCTCTGTTTTGTGGCTGCTGGGGCAGGTGCGACACTGCCGCTTTCTCTAAAGATGCCCCCAAGCTGCTCTGGGGGTAGAAGGAAGGCTGCAGGGAACAGCAGGACCCGGAGAACCCAGACCCCAAGGACCAGGTGGTCCTTAAGCCATCCTGCAGGAGGCCGGCTGCTCTAGCAGGTTGAGGAGTGCCACGCGGCTCTGTGGGTACCGCCTCACTTGGCCTCGGCGGCCAAGCTTAGACACTAGA

## >CaRE4-Scramble

CCTACTGGCCGGTACCTGAGCTCGCTAGCCTCCCTCTTCCTCCTGTACCCTCAACCCATCAGCTGCCTGTACTGCTGTAGCAACAACTTCTCTCTGAAAACCTTCTATTTCTGGTGGCTACACACAGGTGTCTGCTCCGGCAGGATATCTCCCAGAGGTGTTTATAGACTCCGT AAAGAT C GGC AC GAC AGGAGT C AAC AC T GGT GC GGAGAAAC GAAAGC AC GT C AAGGCGGCAGAGAGTGCTACGGTGCGGCTGGCGAGGGGAAGCGTCCTTCTTCAGAATTCTGGCTGTTATCCTCTTACTTGGCACGAGCTCCGATGGGGTCCGTACAAGTGACCGGCCTACGAGGTCGGCACGTAAGGTGGAGTGTCCTTTACTAATAAGCCCGGTTTGATAACAGACCGTGCAGGACCGAGGACTGCACTGGATTTCACCCTAAAGCCCCGTAGCGCCTTAAGCCATCCTGCAGGAGGCCGGCTGCTCTAGCAGGTTGAGGAGTGCCACGCGGCTCTGTGGGTACCGCCTCACTTGGCCTCGGCGGCCAAGCTTAGACACTAGA

## >CaRE4-SNP

CCTACTGGCCGGTACCTGAGCTCGCTAGCCTCCCTCTTCCTCCTGTACCCTCAACCCATCAGCTGCCTGTACTGCTGTAGCAACAACTTCTCTCTGAAAACCTTCCCAGGGGAGAAAACGCGCCCCCCCTCACGCCTCAGAGGCACACACCTATCCTAGCCTTTGTTTTTTTATTGCTGAGAGTACAGAAAGCAGCGGCTTCTGAAGGAGGTATTTATTTTGGTCCCAAAC AGAAAAGAGT GGGT GAT TCTGT GGGC AGAGGT GGC AATT CCT GAAT GT GGT AT GGTGACTCACGCCCGGAGGTGCTCAGAATTTCCTCTGTTTTGTGGCTGCTGGGGCAGGTGCGACACTGCCGCTTTCTCTAAAGATGCCCCCAAGCTGCTCTGGGGGTAGAAGGAAGGCTGCAGGGAACAGCAGGACCCGGAGAACCCAGACCCCAAGGACCAGGTGGTCCTTAAGCCATCCTGCAGGAGGCCGGCTGCTCTAGCAGGTTGAGGAGTGCCACGCGGCTCTGTGGGTACCGCCTCACTTGGCCTCGGCGGCCAAGCTTAGACACTAGA

## Generation of CRISPR Mouse Models

### 12DEL Mouse

12DEL mice were generated by the UCSF Mouse Genetic core (San Francisco, CA, USA) by microinjection of Cas9 ribonucleoprotein (PNA Bio, Newbury Park, CA, USA) into C57BL/6 zygotes. Briefly, Cas9 (50 ng/*μ*L) mIL2RA-CaRE4 gRNA-1 (25 ng/*μ*L), and ssDNA HDR template (50 ng/*μ*L) were mixed in injection buffer (10 mM Tris, 0.1 mM EDTA) and incubated on ice for 10 minutes, as per the manufacturer’s instructions. The mixture was microinjected into the cytoplasm of C57BL/6 single-cell zygotes isolated from super-ovulated females. We did not observe knock-in of the SNP in the progeny, but a single founder carried a 12 bp deletion in the *IL2RA* intronic enhancer. The 12DEL mouse line was established by backcrossing this founder for at least one generation before breeding to homozygosity. gRNA and HDR template sequences are listed in Supplementary Table 3.

### SNP Mouse

SNP knock-in mice were generated by the Jackson Laboratory (Bar Harbor, ME, USA) by microinjection of gRNA and Cas9 mRNA. Briefly, Cas9 mRNA (100 ng/μL), mIL2RA-CaRE4 gRNA-1 (50 ng/μL), and ssDNA HDR template (100 ng/μL) were mixed and injected into C57BL/6 zygotes. Three founders with the knock-in SNP were identified by PCR amplicon sequencing and confirmed by sequencing of TOPO-cloned PCR products. The SNP mouse lines were established by backcrossing founders for at least one generation before breeding to homozygosity. gRNA and HDR template sequences were identical to those used to generate the 12DEL mouse line and are listed in Supplementary Table 2.

## Mouse Genotyping

All mice were genotyped by Sanger sequencing genomic DNA from proteinase K digested tail tissue. PCR amplification of the CaRE4 enhancer was carried out using HotStart Taq (Bioline USA Inc, Taunton, MA, USA) and primers (mIL2RA-CaRE4-F, mIL2RA-CaRE4-R) that span the edited site. PCR amplicons were then sequenced with the mIL2RA-CaRE4-F primer.

## Mouse Immunophenotyping

### Cell preparation

Mice were maintained in the UCSF specific pathogen-free animal facility in accordance with guidelines established by the Institutional Animal Care and Use Committee and Laboratory Animal Resource Center. All mouse experiments were done with littermate controls on animals aged 2–4 months. Spleen, peripheral lymph nodes (peri-LNs), thymus and large intestine was collected from each mouse. Spleen, peri-LNs, and thymus were dissociated in 1x PBS with 2% FBS and 1 mM EDTA. The mixture was then passed through a 70 μm filter. ACK lysis was used to deplete red blood cells from splenocytes.

For the large intestines, we used a lamina propria dissociation kit (Miltenyi Biotec, San Diego, CA, USA; Cat#130–097-410) and a gentleMACS dissociator (Miltenyi Biotec, San Diego, CA, USA) to isolate lamina propria lymphocytes (LPLs) from the large intestine. The LPLs were then enriched for CD4+ T cells using a negative selection kit (StemCell Technologies, Vancouver, BC, CA; Cat#19752). Cell counts were taken using a Coulter Counter.

### Staining

All antibody stains were performed at a 1:100 dilution in 30 μL of 1x PBS. To pellet the cells centrifugation was done at 400xg for 5 minutes. For immunophenotyping, two million cells were stained per tissue sample. Cells were first stained with a viability dye at a 1:1000 dilution in 1x PBS for 20 minutes at 4 °C, then washed with EasySep Buffer (1x PBS, 2% FBS, 1mM EDTA). Cells were then resuspended in the appropriate surface staining antibody cocktail and incubated for 30 minutes at 4 °C, then washed with 1x PBS. Cells were then fixed, permeabilized, and stained for transcription factors using the FOXP3 staining kit (eBioscience, Cat#00–5523-00) according to the manufacturer’s instructions. Antibody staining panels are listed in Supplementary Table 1.

## Primary Human Naïve T cell Editing

### Naïve T cell Isolation

PBMCs were isolated from fresh human whole blood using SepMate tubes (StemCell Technologies, Vancouver, BC, CA; Cat#85450). PBMCs were then enriched for CD4+ T cells using EasySep CD4 negative selection kit (StemCell Technologies, Vancouver, BC, CA; Cat#17952). CD4+ T cells were stained with CD4-PerCP, CD127-PE, CD25-APC, CD45RA-VF450, and CD45RO-FITC. Naïve T cells were sorted as CD4+CD25^low^CD127^high^CD45RA+CD45RO- and rested overnight in RPMI supplemented with 10% FBS, 1 mM sodium pyruvate, non-essential amino acids, 2 mM glutamine, 100 U/ml penicillin, 100 μg/mL streptomycin and 10 mM HEPES.

### Electroporation of Primary Human T cells

T cells were electroporated with Cas9 ribonucleoproteins and homology directed repair (HDR) templates^6^. Inclusion of the ssDNA HDR templates could disrupt the target sequences either through “knock-in” of the template sequence or by increasing indel formation. Briefly, 80 μM crRNA was mixed 1:1 with 80 tracRNA and incubated at 37 °C for 30 minutes to form a gRNA. The gRNA was then mixed with 40 μM Cas9 protein and incubated at 37 °C for 10 minutes to form the Cas9 RNP. During the incubation, 1 million naïve T cells were resuspended in 20 ul of P3 buffer (Lonza, Basel, Switzerland) and transferred to an electroporation well. We then added 6 ul of Cas9 RNP and 100 umol of HDR template. The cells were nucleofected using the Amaxa 4D, program EH115. Cells were then transferred to a round-bottom 96 well plate and immediately activated in complete RPMI with anti-CD3 anti-CD28 dynabeads at a 3:1 bead:cell ratio. FACS analysis and sorting of the cells was done at days 1, 2 and 3 post-activation. IL2RA negative and positive gates were established using unstained cells. gRNA reagents were commercially synthesized (Dharmacon).

## Amplicon sequencing

Genomic DNA was extracted from sorted cells using QuickExtract (Epicentre). PCR amplicons from the edited regions were prepared for next-generation sequencing using a nested PCR strategy. The outer PCR (PCR1) generates a 600–700 bp amplicon using primers outside the region matching the HDR donor template. The inner PCR (PCR2) generates the ~250 bp amplicon to be sequenced. Primer sequences are listed in Supplementary Table 2.

For PCR1, genomic DNA from 5,000–10,000 sorted cells was amplified using PrimeStar GXL DNA Polymerase (Clontech, Cat. No. R050B) according to the following protocol: 98 °C, 2’; 30 cycles at 98 °C, 10”, 55 °C, 15”, 68 °C, 45”. PCR1 products were purified using SPRI beads and quantified using the Qubit dsDNA HS Assay Kit (Thermo). The purified products of PCR1 (20 ng/sample) were used as templates for PCR2 and amplified using PrimeStar GXL DNA Polymerase according to the following protocol: 98 °C, 2’; 10 cycles at 98 °C, 10”, 60 °C, 15”, 68 °C, 20”. PCR2 products were purified and quantified as done for PCR1. Indexed sequencing libraries were prepared with the TruSeq Nano DNA HT Library Prep Kit (Illumina, Cat. No. FC121–4001) using 100 ng of purified PCR2 product. Libraries were pooled and 150 bp paired-end sequencing was performed with a MiSeq sequencer (Illumina).

## Amplicon-seq analysis

Fastq files for paired end sequencing of the *IL2RA* exon and enhancer edited sites were processed using Trimmomatic^8^ (leading: 15, trailing: 15, headcrop: 17, sliding window:4:15). The trimmed fastq files were then aligned to the human reference genome (build hg19/GRCh37) using the Burrows Wheeler Aligner (BWA-MEM) algorithm^9^. The samtools mpileup utility was used to quantify the total number of reads that mapped to each position surrounding the cut sites, and a custom script examining the CIGAR string was used to estimate the number and locations of insertions and deletions for each read. Reference sequence enrichment scores at individual bases were calculated by dividing the percent reference sequence at a given base in sorted IL2RA- or IL2RA+ cells by the percent reference sequence at the same position in unsorted edited cells.

## SNP Knock-in Analysis

For human primary T cell editing of the IL2RA enhancer we included HDR templates with the rs61839660 SNP or a coding mutation (c.530G->A) to introduce a premature stop codon. The repair template was added primarily to boost editing efficiency of the Cas9 RNPs. This also provided us with the opportunity to assess the effect of the SNP on IL2RA expression in isogenic primary human T cells by following SNP knock-in alleles in IL2RA- and IL2RA+ cells with amplicon sequencing. Efficient T cell genome editing and HDR is only possible in activated cells. Here we activated the cells immediately after electroporation with Cas9 RNP and the HDR template. We noticed that both the number and size of insertion and deletion (indel) edits increased over the three-day time course consistent with continuous Cas9 cutting and/or imperfect DNA repair activity for the duration of the experiment. Similarly, HDR increased over the three days following electroporation and stimulation. We observed relatively few SNP knock-in alleles at day 1, increasing 20-fold by day 3 (up to ~2% of total alleles) (Supplementary Fig. 9). Despite the dynamic editing of the cells due to the stimulation requirement, we were able to normalize edits in the IL2RA- and IL2RA+ cells by comparing to the bulk population at each time point. Enrichment of SNP knock-in in the IL2RA-population is consistent with the SNP impairing IL2RA expression. When we looked at the distribution of HDR alleles we observed enrichment of the SNP in the IL2RA-population in both donors, for all three days (Supplementary Fig. 9). Tracking variant knock-in alleles may be a generalizable approach to studying how variants affect gene expression in an isogenic background.

